# NMDA receptors enhance the fidelity of synaptic integration

**DOI:** 10.1101/566117

**Authors:** Chenguang Li, Allan T. Gulledge

## Abstract

Excitatory synaptic transmission in many neurons is mediated by two co-expressed ionotropic glutamate receptor subtypes, AMPA and NMDA receptors, that differ in their kinetics, ion-selectivity, and voltage-sensitivity. AMPA receptors have fast kinetics and are voltage-insensitive, while NMDA receptors have slower kinetics and increased conductance at depolarized membrane potentials. Here we report that the voltage-dependency and kinetics of NMDA receptors act synergistically to stabilize synaptic integration of excitatory postsynaptic potentials (EPSPs) across spatial and voltage domains. Simulations of synaptic integration in simplified and morphologically realistic dendritic trees revealed that the combined presence of AMPA and NMDA conductances reduces the variability of somatic responses to spatiotemporal patterns of excitatory synaptic input presented at different initial membrane potentials and/or in different dendritic domains. This moderating effect of the NMDA conductance on synaptic integration was robust across a wide range of AMPA-to-NMDA ratios, and results from synergistic interaction of NMDA kinetics (which reduces variability across membrane potential) and voltage-dependence (which favors stabilization across dendritic location). When combined with AMPA conductance, the NMDA conductance balances voltage- and impedance-dependent changes in synaptic driving force, and distance-dependent attenuation of synaptic potentials arriving at the axon, to increase the fidelity of synaptic integration and EPSP-spike coupling across neuron state (i.e., initial membrane potential) and dendritic location of synaptic input. Thus, synaptic NMDA receptors convey advantages for synaptic integration that are independent of, but fully compatible with, their importance for coincidence detection and synaptic plasticity.

**Significance Statement:** Glutamate is an excitatory neurotransmitter that, at many synapses, gates two coexpressed receptor subtypes (AMPA and NMDA receptors). Computational simulations reveal that the combined synaptic presence of AMPA and NMDA receptors reduces variability in synaptic integration in response to identical patterns of synaptic input delivered to different dendritic locations and/or at different initial membrane potentials. This results from synergistic interaction of the slower kinetics and voltage-dependence of NMDA receptors, which combine to enhance synaptic currents when synaptic driving forces are otherwise reduced (e.g., at depolarized membrane potentials or in distal, high-impedance dendrites). By stabilizing synaptic integration across dendritic location and initial membrane potential, NMDA receptors provide advantages independent of, but fully compatible with, their well-known contribution to synaptic plasticity.

## Introduction

In the vertebrate central nervous system, fast excitatory synaptic transmission is mediated primarily by the amino acid glutamate, which at many synapses gates two coexpressed ionotropic receptors: α-amino-3-hydroxy-5-methyl-4-isoxazolepropionic acid (AMPA) and N-methyl-D-aspartate (NMDA) receptors. While both receptor subtypes are gated by glutamate, and are permeable to monovalent cations, NMDA receptors are distinct in having slower kinetics, permeability for calcium, and voltage-dependence due to channel blockade by extracellular magnesium ions at hyperpolarized membrane potentials (for review, see Iacobucci and Popescu, 2017). These features of NMDA receptors, which are highly conserved across phyla (Greer et al., 2017), are critical for their well-established role in gating associative synaptic plasticity, including certain types of long-term potentiation and depression (for review, see Luscher and Malenka, 2012). NMDA receptors also influence synaptic integration, as the voltage-dependence of NMDA receptors can promote linear (Cash and Yuste, 1998) or supralinear (Schiller et al., 2000) summation of excitatory postsynaptic potentials (EPSPs) that occur in sufficient spatial and temporal proximity to relieve magnesium block of NMDA channels, and can reduce the voltage-dependent variability of mixed AMPA/NMDA synaptic currents (Diamond and Copenhagen, 1993; Connelly et al., 2016).

Because the input impedance of dendrites increases with distance from the soma, distal synapses generate larger, more depolarizing local dendritic EPSPs than do proximal synapses (Jaffe and Carnevale, 1999; Gulledge et al., 2005; Lajeunesse et al., 2013). Yet the ability of an individual AMPA-mediated EPSP to recruit synaptic NMDA conductance is tempered by its rapid kinetics (∼0.2 and 2.0 ms activation and decay, respectively), which, when combined with the limited local capacitance of narrow dendrites and dendritic spines, generates EPSPs that decay too rapidly to efficiently recruit slower activating (> 2 ms) NMDA conductances (Stern et al., 1992; Gulledge et al., 2012). Instead, network-driven patterns of synaptic input interact within the dendritic tree based on their spatiotemporal relationships to recruit NMDA conductances (Polsky et al., 2009; Grienberger et al., 2014; Palmer et al., 2014; Cichon and Gan, 2015), especially when they occur in high-impedance dendritic branches or spines (Branco and Hausser, 2011; Gulledge et al., 2012; Harnett et al., 2012). NMDA conductances are also recruited when barrages involve repetitive activation of the same synapses, as the extended occupancy of glutamate sites on NMDA receptors (∼100 ms) “prime” them for immediate gating during subsequent AMPA-mediated local depolarization (Schiller et al., 2000; Polsky et al., 2004). With sufficient levels of synaptic activation, inward NMDA currents become self-sustaining “NMDA spikes” that amplify and prolong synaptic depolarization to generate supralinear summative events at the soma and axon (for reviews, see Antic et al., 2010; Branco and Hausser, 2010; Major et al., 2013; Grienberger et al., 2015).

While most prior studies examining the impact of synaptic conductance on the location- and/or voltage-dependence of synaptic transmission have focused on single synaptic events (e.g., Jaffe and Carnevale, 1999; Lajeunesse et al., 2013), we reasoned that the voltage-dependency of NMDA receptors, combined with the distance-dependent electrotonic structure of dendritic trees, should reduce the variability of somatic drive (as measured as somatic depolarization and/or action potential generation) in response to spatiotemporal patterns of synaptic input occurring at different dendritic locations and/or from different initial membrane potential states. Here we test this hypothesis using computational simulations to compare the impact of synaptic conductance (AMPA-only, NMDA-only, or both conductances together) on the integration of patterns of afferent input delivered to different dendritic locations and/ or at different initial membrane potentials. Our results reveal that AMPA and NMDA conductances interact in a complementary fashion to stabilize synaptic integration and EPSP-spike coupling in ways that are independent of, yet fully compatible with, their well-established roles in coincidence detection and synaptic plasticity.

## Materials and Methods

### Computational models

Simulations were made using NEURON 7.7 software (Carnevale and Hines, 2006; RRID: SCR_005393) and the Neuroscience Gateway portal (Sivagnanam et al., 2013). The code used to generate the data in this paper is available online at ModelDB (http://modeldb.yale.edu/266802; access code = “nmda”). Morphologies used include “ball-and-stick” model neurons, consisting of somata connected to spinous dendrites of variable length, a hippocampal CA3 pyramidal neuron (Hemond et al., 2009), and a hippocampal dentate granule neuron (Schmidt-Hieber et al., 2007). **Table 1** lists the dimensions and membrane parameters for all neuron morphologies. Active mechanisms consisted primarily of fast-inactivating voltage-gated sodium and delayed-rectifier potassium conductances (source codes available in <u>ModelDB</u>, entry <u>144385</u>). Spines with neck resistances of 500 MΩ (Harnett et al., 2012) were positioned at 1-µm intervals along dendrites. Axons in all models were 2,000 µm long, had diameters of 0.5 µm, and were attached to the soma via a 40 µm axon initial segment (AIS) that tapered from 2.0 µm (at the soma) to 0.5 µm (at the axon). Except for the AIS, voltage-gated conductances were evenly distributed in axons (i.e., axons were modeled as unmyelinated; see **Table 1** for sodium and potassium channel densities in each neuronal compartment). Unless otherwise noted, ball-and-stick neurons included somata (20 ⨯ 10 µm) attached to a single tapering (5 µm to 1 µm) dendrite of variable length (0.2 to 1 mm; input resistances (R_N_) ranging from ∼125 to ∼310 MΩ). Because R_N_ can influence the impact of dendritic location on synaptic integration, for some simulations we delivered synaptic input to a 200-µm dendrite attached to a larger neuron typically having three additional 600-µm dendrites that did not receive synaptic input (**Figure 3A** shows results from a range of neurons having from 1 to 5 additional 600-µm dendrites). In all simulations, models were initiated following a 1 s passive run to allow active conductances to reach a steady-state. Simulations were run with time steps of 10 or 25 µs at a nominal temperature of 37°C.

**Table 1:** Model parameters.

| Neuron Morphology | Compartment | Dimensions (l x w) | # of Segments | Active properties (max. conductance) |
| --- | --- | --- | --- | --- |
| Ball-and-stick* | Soma | 20 x 10 $\mu\text{m}$ | 3 | Na<sup>+</sup>: 100 pS/$\mu\text{m}^2$<br>K<sup>+</sup>: 100 pS/$\mu\text{m}^2$ |
| | Dendrite | 200 - 1,000 $\mu\text{m}$, tapering from 5 to 1 $\mu\text{m}$ | 1 per $\mu\text{m}$ | Typically passive.<br>Figure 3C: Na<sup>+</sup> and K<sup>+</sup> with linear decrease (100 to 10 pS/$\mu\text{m}^2$). |
| CA3* and dentate granule neurons | Soma | As reconstructed | 3 | Na<sup>+</sup>: 100 pS/$\mu\text{m}^2$<br>K<sup>+</sup>: 100 pS/$\mu\text{m}^2$ |
| | Dendrites | As reconstructed | 1 per $\mu\text{m}$ | Passive. |
| All morphologies | Spines | Neck: 1 x ~0.05 $\mu\text{m}$<br>Head: 0.5 x 0.5 $\mu\text{m}$ | 1<br>1 | As in parent dendritic compartment. |
| | Axon initial segment | 40 $\mu\text{m}$, tapering from 2 to 0.5 $\mu\text{m}$ | 40 | Na<sup>+</sup>: 100 pS/$\mu\text{m}^2$ (1<sup>st</sup> 5 $\mu\text{m}$) or 8000 pS/$\mu\text{m}^2$<br>K<sup>+</sup>: 100 pS/$\mu\text{m}^2$ (1<sup>st</sup> 5 $\mu\text{m}$) or 2000 pS/$\mu\text{m}^2$ |
| | Axon | 2000 x 0.5 $\mu\text{m}$ | 201 | Na<sup>+</sup>: 300 pS/$\mu\text{m}^2$<br>K<sup>+</sup>: 60 pS/$\mu\text{m}^2$ |

\*The ball-and-stick model in Figure 1 and the CA3 neuron in Figure 8 are purely passive. General model parameters: $R_M = 15 \text{ k}\Omega\cdot\text{cm}^2$ ; $C_M = 1 \text{ }\mu\text{F}/\text{cm}^2$ ; $R_i = 100 \text{ }\Omega\cdot\text{cm}$ ; $E_{\text{pas}} = -55 \text{ to } -85 \text{ mV}$ , as indicated in text; time steps, 10 to 25 $\mu\text{s}$ ; nominal temperature, 37°C.

**Figure 1.**
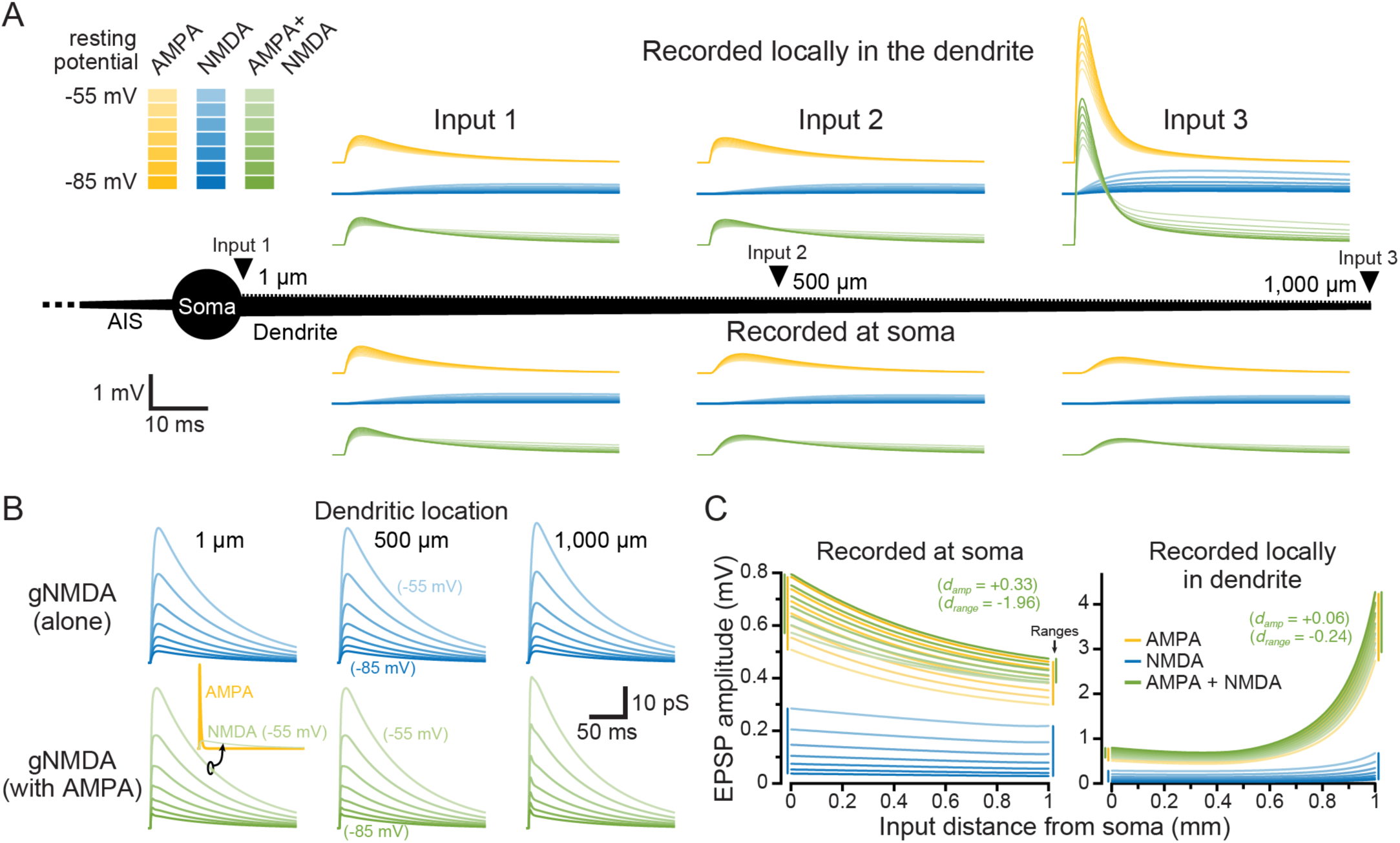
Impact of membrane potential and dendritic location on synaptic conductances and their resulting EPSPs. ***A***, Individual synaptic inputs were activated along the dendrite (1,000 µm length) of a passive ball-and-stick model neuron (black; not to scale). Shown are somatic (bottom) and local dendritic (top) EPSPs generated by synapses placed at three dendritic locations (1, 500, and 1,000 µm from the soma), at seven different resting membrane potentials (RMPs; -55 to -85 mV, as indicated by color depth). AMPA-only EPSPs are shown in yellow, NMDA-only synapses are shown in blue, and inputs having both AMPA and NMDA conductance are shown in green. ***B***, The NMDA conductances underlying the EPSPs shown in ***A***. Inset are the AMPA (yellow) and NMDA (green) conductances for the AMPA+NMDA input at -55 mV at the most proximal dendritic location (shown to scale with the 500 pS AMPA conductance). ***C***, Plots of the amplitudes of EPSPs having the indicated synaptic conductances measured at the soma (left) or locally at the site of synaptic input (right), verses dendritic location of the synapse. Synaptic conductances are color-coded as in ***A*** for seven RMPs from -85 to -55 mV, at 5 mV intervals. Inset are the effect sizes (*d*) for AMPA+NMDA EPSP amplitudes (across all dendritic locations and RMPs) and ranges (across all RMPs) relative to AMPA-only EPSPs. Colored vertical bars at margins indicate the ranges of amplitudes across all RMPs for the most proximal and most distal inputs.

**Figure 2.**
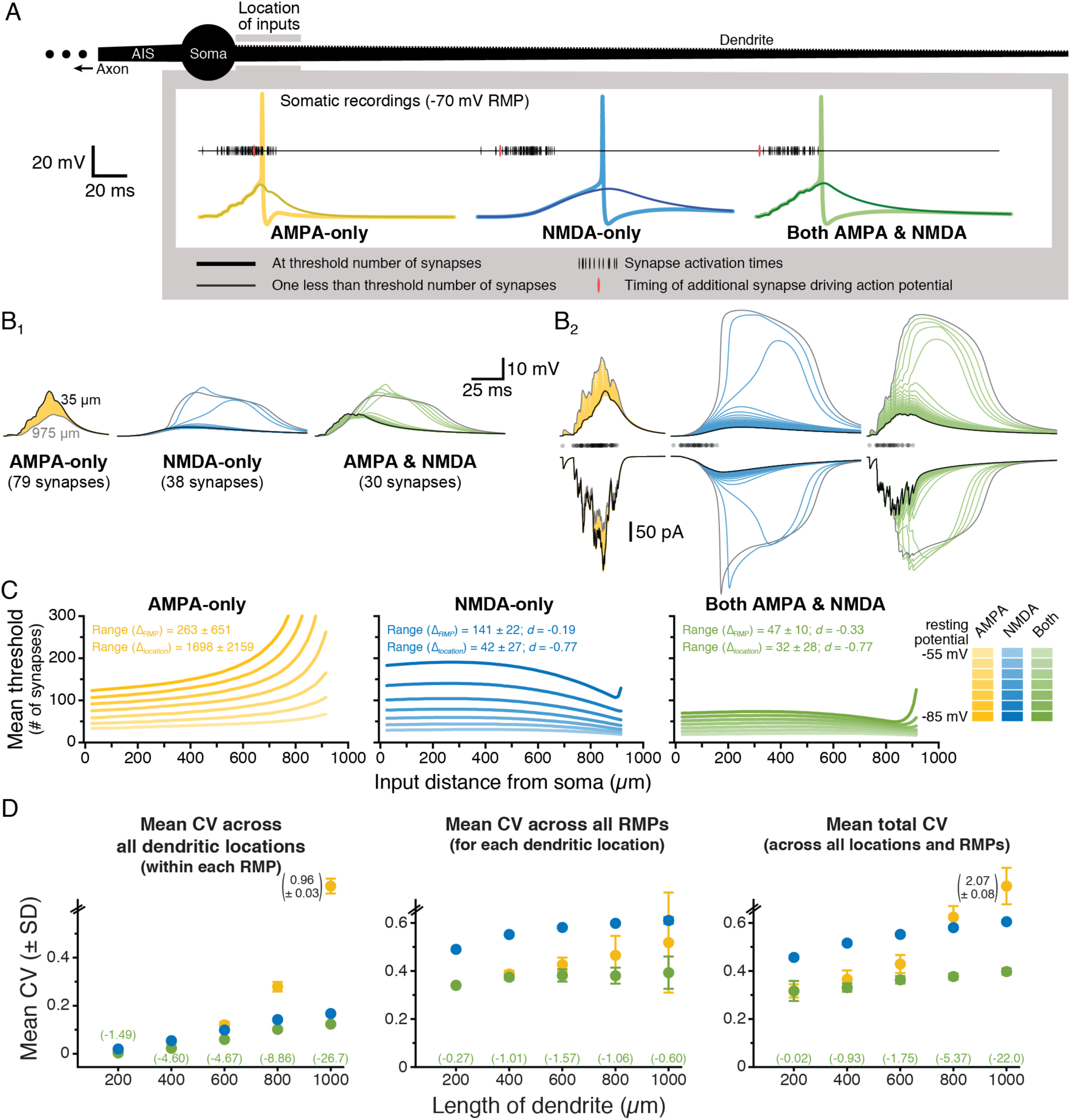
Combining synaptic AMPA and NMDA conductances stabilizes EPSP-spike coupling. ***A***, A spiny ball-and-stick neuron (black) receiving progressively longer iterative trains of a spatiotemporal pattern of synaptic input. Just-threshold (thick traces) and just-subthreshold (thin traces) voltage responses in the soma for AMPA-only (yellow), NMDA-only (blue), or both AMPA and NMDA (green) synaptic conductances in response to inputs arriving within the first 50 µm of the dendrite. Resting membrane potential (RMP) is -70 mV. **B**, Somatic (**B**_**1**_) and dendritic (**B**_**2**_, top) voltage responses for identical subthreshold synaptic barrages at each dendritic location (superim-posed; 20 µm intervals centered between 35 µm [black traces] and 975 µm [gray traces] from the soma) for the indicated synaptic conductance types (RMP is -70 mV). Summed total synaptic currents are shown in **B**_**2**_, bottom. Timings of synaptic activations are shown with semi-transparent black dots above the synaptic currents in **B**_**2**_. ***C***, Plots of the mean threshold numbers of synaptic activations necessary to initiate action potentials at different locations in the dendrite for synaptic inputs having AMPA-only (left; yellow), NMDA-only (middle; blue), or AMPA and NMDA (right; green) conductances, across seven different RMPs (−85 to -55 mV), as indicated by color depth. Mean ranges of synaptic thresholds measured across RMP (Δ_RMP_) or dendritic location (Δ_location_; ± standard deviations) and effect sizes (in units of *s*_AMPA-only_) of conductance on ranges (for NMDA-only and AMPA+NMDA inputs) are inset. ***D***, Plots of mean CVs (± standard deviations) calculated for the threshold number of synapses for each pattern of input (n = 10), across all locations within each RMPs (left), across RMP for each dendritic location (middle), and across all RMPs and dendritic locations (“total CV”; right) for inputs having AMPA-only (yellow), NMDA-only (blue), or both AMPA and NMDA (green) conductances in dendrites of the indicated lengths. Note that in the 1,000 µm dendrite the CV for AMPA-only responses became very large (off scale). The magnitudes of these large CVs is indicated next to their symbol (standard deviation bars are to scale). The effect sizes (*d*) for changes in CV with AMPA+NMDA inputs (relative to AMPA-only CVs; expressed in units of *s*_AMPA-only_) are shown in green for each dendritic length.

**Figure 3.**
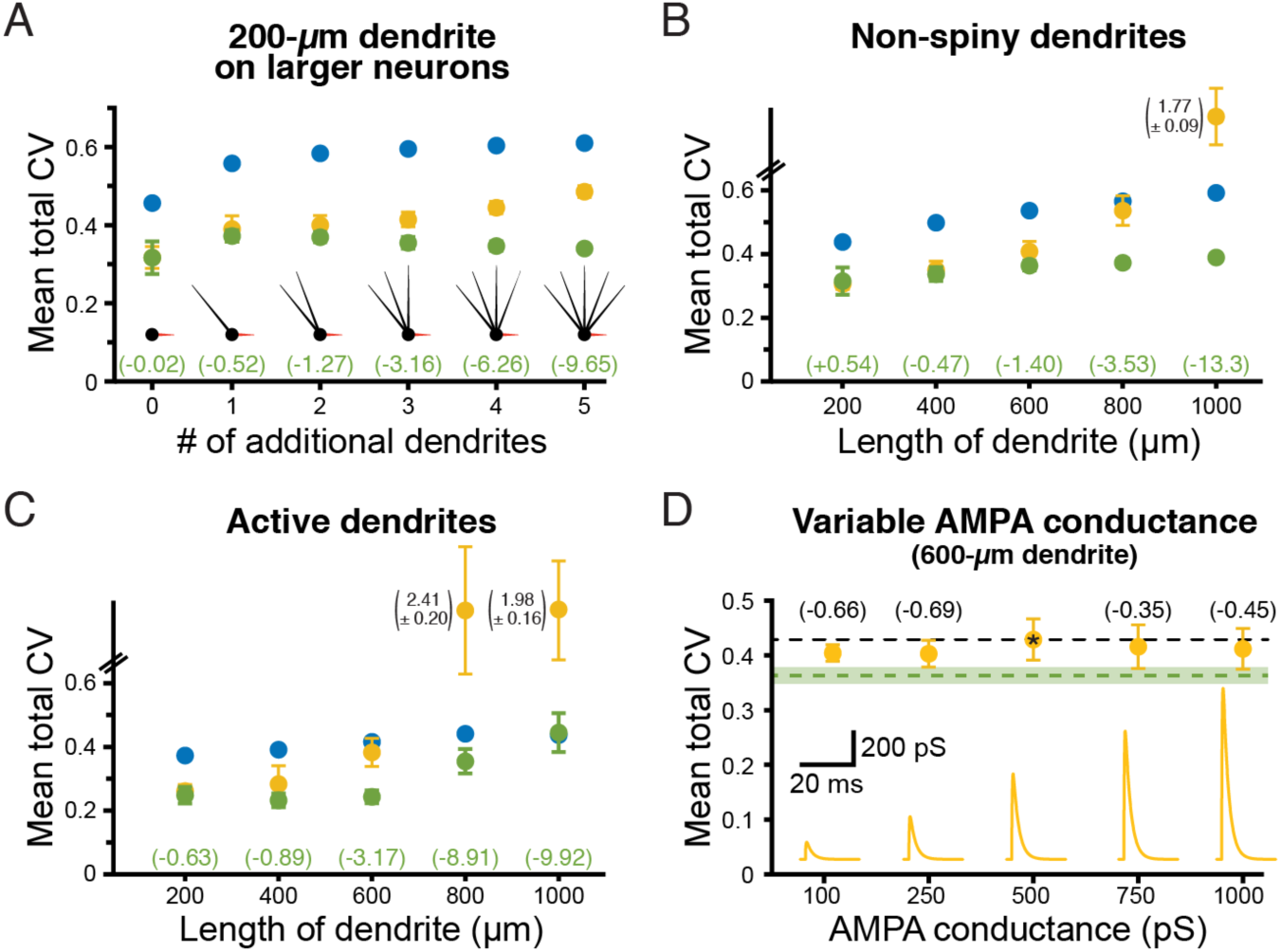
NMDA-dependent stabilization of EPSP-spike coupling is consistent across a range of model conditions and is not dependent on total synaptic conductance. ***A***, Total CVs (± standard deviations) for synaptic thresholds measured across all dendritic locations and RMPs for AMPA-only (yellow), NMDA-only (blue), and AMPA+NMDA (green) synaptic inputs to a 200-µm dendrite placed on a neuron having 0 to 5 additional dendrites (600 µm each, as indicated in inset diagrams). Effect sizes for AMPA+NMDA inputs (relative to AMPA-only inputs) are listed in green (in units of *s*_AMPA-only_). The neuron with three additional dendrites is used in later figures and referred to as the “200-µm dendrite on a larger neuron.” ***B***, Total CVs (± standard deviations) for synaptic thresholds for AMPA-only (yellow), NMDA-only (blue) and AMPA+NMDA (green) inputs to non-spiny dendrites of the indicated lengths. Effect sizes for AMPA+NMDA CVs shown in green (units of *s*_AMPA-only_). ***C***, Total CVs (± standard deviations) for synaptic thresholds measured across all dendritic locations and RMPs for inputs onto spiny dendrites with active conductances (see **Table 1**). ***D***, Plot of mean total CVs for synaptic thresholds (± standard deviations) for AMPA-only EPSPs with the indicated peak conductance magnitudes in a 600 µm dendrite. Black dashed line indicates the mean CV of the standard 500 pS AMPA conductance (indicated by asterisk), while the green dashed line indicates mean total CV (with shaded standard deviation) for AMPA (500 pS) + NMDA (1 nS) inputs in the same dendrite. Effect sizes of peak conductance manipulations, relative to the 500 pS AMPA input, are shown in black (in units of *s*_500pS_). Inset are the AMPA conductances to scale.

### Simulated synaptic inputs

In most simulations, synaptic conductances were located on spine heads. AMPA conductances (ModelDB entry <u>120798</u>) typically had exponential rise and decay time constants of 0.2 ms and 2 ms, respectively, a reversal potential of 0 mV, and a maximum conductance of 500 pS, unless otherwise noted. NMDA inputs were based on the AMPA conductance, with added voltage-dependence (as in Model DB entry <u>184725</u>), and unless otherwise noted had nominal maximum conductances of 1 nS (typically reaching ∼46 pS during individual AMPA+NMDA synaptic events occurring from a membrane potential of -55 mV; see also **Figures 1B** and **4A**), reversal potentials of +5 mV, and exponential rise and decay time constants of 3 ms and 90 ms, respectively. Single synaptic potentials were monitored at the soma and at the dendritic site of synaptic input. For barrages of synaptic input, dendritic voltage responses were measured at the soma and at the mid-point of the dendritic span receiving synaptic input. During spike threshold tests, synaptic thresholds were determined by monitoring action potentials in the midpoint of the axon. During non-spike threshold tests, synaptic thresholds were determined as somatic events reaching 5 mV (for the dentate granule cell) or 3 mV (for the CA3 neuron) above the resting membrane potential (RMP).

Synaptic barrages of *n* synaptic activations, delivered to discrete dendritic compartments, were defined by sampling two random variables *n* times for a set of *n* pairs. The first random variable in the pair determined the site of the *n*^*th*^ synaptic input and was chosen from a uniform distribution spanning integers 0 to 49 (for a 50-µm range), which was then applied to the dendritic segment of interest (i.e., from 620-669 µm from the soma). The second random variable, which determined synaptic timing, was chosen from a Gaussian distribution (width of 50 ms). Each pattern maintained a single ordered set of sampled pairs. The size of the set varied depending on the number *n*, but the content and order of the pairs did not change for a given pattern. For instance, within one pattern, barrages with *n* and *n+1* synaptic activations were identical except for the *n+1*^*th*^ additional activation. These spatiotemporal patterns of synaptic input were iteratively moved along dendrites at 10- or 20-µm intervals to compare the effect of input location on local and somatic synaptic responses. Ten different randomized spatiotemporal patterns of synaptic input were utilized to allow statistical determination of the stability of EPSP-spike coupling over input pattern, dendritic location, and RMP. Nominal RMPs were set by uniformly adjusting the reversal potential for the passive leak conductance in all model compartments.

Variation of EPSP-spike coupling and somatic drive was quantified as coefficient of variation (CV; standard deviation normalized by mean), which provides a relative measure of variability in synaptic thresholds across all dendritic locations and/or membrane potentials independent of response magnitudes (as opposed to measures of absolute range or standard deviation, which are difficult to compare across models having different mean thresholds, and which may overestimate effective variability if functions [e.g., vs distance or voltage] are non-linear). In the longest dendrites tested (the 1,000-µm ball-and-stick and CA3 neurons), inputs to distal dendritic locations became so remote that threshold numbers of synaptic inputs rose exponentially to infinite values (i.e., beyond our maximum tested value of 10,000), as at very distal distances even voltage-clamping a 50-µm dendritic segment to 0 mV would fail to generate action potentials in the axon. Trials with AMPA-only inputs always hit this limit earliest (i.e., at slightly more proximal dendritic locations than other conductances), and so CVs for all conductances (AMPA-only, NMDA-only, or both AMPA and NMDA) were calculated only across dendritic locations where AMPA synaptic thresholds were measurable.

### Statistical analyses

Results are reported as means ± standard deviations. Because our simulations are deterministic, we know *a priori* that differences in synaptic responses are “real” (that is to say, we know they result from the changes in synaptic conductance coded into the simulation). To quantify the relative impact of synaptic conductance on variability of synaptic integration (as measured by CV of synaptic thresholds) across RMPs and/or dendritic locations, we calculated the effect size (*d*) for each synaptic conductance as the mean difference in CV (relative to AMPA-only trials) normalized by the standard deviation (*s*) of the AMPA-only trials, according to the following formula:

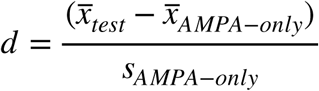

where 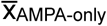 and 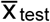 are the means of CVs of synaptic thresholds calculated across trials (i.e., ten patterns of synaptic input) for AMPA-only simulations and those with modified synaptic conductances, respectively, and *s*_AMPA-only_ is the standard deviation for CVs in AMPA-only trials. Thus, *d* represents both the direction (by sign) and magnitude (in units of *s*_AMPA-only_) of differences in mean variability of synaptic thresholds. This generates a more conservative measure of effect size than other approaches (such as Cohen’s *d*) because the standard deviation of synaptic thresholds for AMPA+NMDA trials was typically much smaller than for AMPA-only trials (see, for instance, the error bars in **Figures 2D** and **6C**). Quantifying effect size in this manner allows direct comparison of the relative impact of synaptic conductance on variability in neurons of different sizes and across different RMPs.

## Results

### Effect of synaptic conductance type on individual EPSPs

We first compared the relative efficacy of individual synapses consisting of AMPA-only, NMDA-only, or both (AMPA and NMDA) conductances in depolarizing the soma of a simplified and fully passive ball-and-stick model neuron (see Methods) with the resting membrane potential (RMP) set to one of seven values (−85 to -55 mV; see Methods). As expected, given the cable properties of dendrites (Gulledge et al., 2005), the amplitude and kinetics of the resulting EPSPs depended on synaptic conductance type, RMP, and the dendritic location of synaptic input (**Figure 1A**). NMDA conductances were larger at depolarized RMPs and/or in distal dendritic compartments (**Figure 1B**), generating NMDA-mediated EPSPs that were largest under those conditions (light-blue plots in **Figure 1C**). On the other hand, AMPA-only and AMPA+NMDA EPSPs had maximal amplitudes at the most hyperpolarized RMPs, where driving force was greatest (dark yellow plots in **Figure 1C**). Across all dendritic locations and RMPs, somatic EPSP amplitudes averaged 0.48 ± 0.10, 0.11 ± 0.07, and 0.52 ± 0.09 mV for AMPA-only, NMDA-only, and AMPA+NMDA responses, respectively. Somatic EPSPs generated by AMPA+NMDA conductances were less variable across dendritic location and RMP (CV = 0.18; effect size, *d*, of +0.33 vs AMPA-only inputs) than EPSPs generated by AMPA-only (CV = 0.22) or NMDA-only (CV = 0.65) inputs. Similarly, AMPA+NMDA somatic EPSPs had smaller absolute amplitude ranges across RMPs for each dendritic location (mean range of 0.14 ± 0.04 mV; *d* = -1.96 relative to AMPA-only inputs) relative to ranges for AMPA-only (0.21 ± 0.03) or NMDA-only (0.21 ± 0.02 mV) EPSPs. While somatic EPSP amplitudes decremented with synaptic distance from the soma, amplitudes at the site of synaptic input became progressively larger at distal dendritic locations (**Figure 1C**), and averaged 0.96 ± 0.68, 0.15 ± 0.11, and 1.00 ± 0.69 mV for AMPA-only, NMDA-only, and AMPA+NMDA inputs, respectively. AMPA+NMDA dendritic EPSPs were marginally larger than AMPA-only EPSPs (*d* = +0.06) and had slightly smaller absolute ranges across RMPs (*d* = -0.24). These data demonstrate that adding an NMDA conductance to individual AMPA-only EPSPs generates marginally larger EPSPs with slightly reduced variability across dendritic location and RMP.

### Impact of synaptic conductance on EPSP-spike coupling

Although the impact of the NMDA conductance on somatic EPSP variability was small for individual EPSPs (compare yellow and green traces in **Figure 1A**), we reasoned that its impact might be amplified when dendrites experience barrages of excitatory synaptic input. To test this, we measured EPSP-spike coupling in ball-and-stick model neurons and compared, across dendritic location and RMP, the threshold number of synaptic activations necessary for action potential initiation in the axon (**Figure 2A**). Ten stochastic spatiotemporal patterns of synaptic input (“synaptic barrages,” see Methods) were delivered to progressively more distal 50-µm-spans of the dendrite. For simulations involving AMPA-only, NMDA-only, or both synaptic conductances, the number of activated synapses within each barrage was iteratively increased until an action potential was initiated. For each pattern of synaptic input, the threshold number of synapses necessary for action potential generation was determined for inputs occurring at different dendritic locations (incremented at 10-µm intervals) and across seven RMPs (−85 to -55 mV). As with single EPSPs, identical barrages of synaptic input generated ever-larger local dendritic depolarization at progressively more distal dendritic locations. Whereas the increase in local response amplitude with distance was linear when inputs involved AMPA conductances only, incorporation of the NMDA conductance, by itself or in combination with the AMPA conductance, allowed for supralinear distance-dependent increases in response amplitude and width, including long-lasting regenerative NMDA spikes, at distal dendritic locations (**Figure 2B**).

For AMPA-only synapses, the mean threshold number of synaptic inputs increased with distance from the soma or with hyperpolarization of the RMP (**Figure 2C**, left). This result reflects 1) distance-dependent voltage attenuation of summated EPSPs as they transfer to the soma, 2) smaller synaptic currents occurring when driving force is reduced during EPSPs in narrow, high-impedance dendrites (where local EPSP amplitudes are intrinsically larger and bring the membrane potential closer to the synaptic reversal potential), and 3) the necessity for larger EPSPs to reach action potential threshold from hyperpolarized RMPs. On the other hand, when synapses contained only the NMDA conductance, the threshold number of synapses decreased with depolarization of the RMP, but was fairly uniform across distance in the proximal dendrite before becoming *lower* at distal locations (**Figure 2C**, middle). This shape resulted from distance-dependent voltage attenuation of somatic EPSPs combined with progressively larger synaptic currents at more distal, high-impedance dendritic locations, where NMDA spikes of increasing amplitude and duration were observed (see **Figure 2B**). Remarkably, when synapses contained both AMPA and NMDA conductances, synaptic thresholds for action potential generation were less sensitive to changes in RMP and less variable across distance than were thresholds for AMPA-only or NMDA-only synaptic inputs (**Figure 2C**). While this reduced variability was evident from the mean ranges of synaptic thresholds (**Figure 2C**), they were just as robust when threshold variability was normalized as CV. Whether measured across all dendritic locations (within any given RMP; **Figure 2D**, left), all RMPs (at any given dendritic location; **Figure 2D**, middle), or across all dendritic locations and RMPs (“total CV”; **Figure 2D**, right), CVs of synaptic thresholds were always lowest when synapses contained both AMPA and NMDA conductances. Effect sizes were amplified in longer dendrites, likely reflecting their greater electrotonic diversity across input locations (see also **Figure 3**, below).

Total CVs for synaptic thresholds, measured across all dendritic locations and RMPs for AMPA-only trials, ranged from a low of 0.32 ± 0.03 (in the 200 µm dendrite) to 2.07 ± 0.08 (in the 1,000 µm dendrite; n = 10 patterns of synaptic input). When synapses incorporated both AMPA and NMDA conductances, CVs ranged from 0.32 ± 0.04 to 0.40 ± 0.01 (in the 200 µm and 1,000 µm dendrites, respectively), with effect sizes (relative to AMPA-only trials) ranging from negligible (−0.02 *s*_AMPA-only_; see Methods) in the 200-µm dendrite, to enormous (−22 *s*_AMPA-only_) in the 1,000-µm dendrite. Synaptic thresholds for NMDA-only conductances were always more variable than for inputs having both conductances, with total CVs ranging from 0.46 ± 0.01 to 0.61 ± 0.01 (in dendrites of 200 and 1,000 µm, respectively) and effect sizes ranging from +3.28 *s*_AMPA-only_ (in the 600-µm dendrite) to -19.3 *s*_AMPA-only_ (in the 1,000-µm dendrite; **Figure 2D**).

To test whether the impact of dendritic length on synaptic threshold variability was reflective of electrotonic structure of the neuron, we measured synaptic thresholds in a 200-µm dendrite attached to larger neurons having 1-to-5 additional dendrites (600 µm each; **Figure 3A**). Even as the measured dendrite remained static (at 200 µm), the effect size of combining AMPA and NMDA conductances on synaptic threshold variability was enhanced as neurons became progressively larger, ranging from almost nothing (−0.02 *s*_AMPA-only_) in the smallest neuron having no additional dendrites, to very large (−9.65 *s*_AMPA-only_) in the largest neuron with five additional dendrites (**Figure 3A**). Similarly, making our normal ball-and-stick neurons electrotonically “tighter” by removing dendritic spines moderately reduced the impact of synaptic conductance on threshold variability (**Figure 3B**).

To test whether the impact of NMDA conductance depended on passive dendrites, in some simulations we added dendritic voltage-gated sodium and potassium conductances (see **Table 1**). Combining AMPA and NMDA conductances continued to reduce the variability of synaptic thresholds relative to AMPA-only inputs when dendrites contained active conductances (**Figure 3C**).

Finally, to test whether the reduced variability in synaptic thresholds for AMPA+NMDA inputs was due to their larger peak conductance magnitudes, we repeated simulations using a range of AMPA-only conductances (100 pS to 1 nS) in a 600-µm dendrite. We found no consistent relationship between peak AMPA conductance and total CV of EPSP-spike coupling measured across all dendritic locations and RMPs (**Figure 3D**; see also **Figure 7**, below). This suggests that, rather than reflecting larger total synaptic conductance, the enhanced fidelity of EPSP-spike coupling observed with AMPA+NMDA inputs depends instead on distinct properties intrinsic to the NMDA conductance.

**Figure 4.**
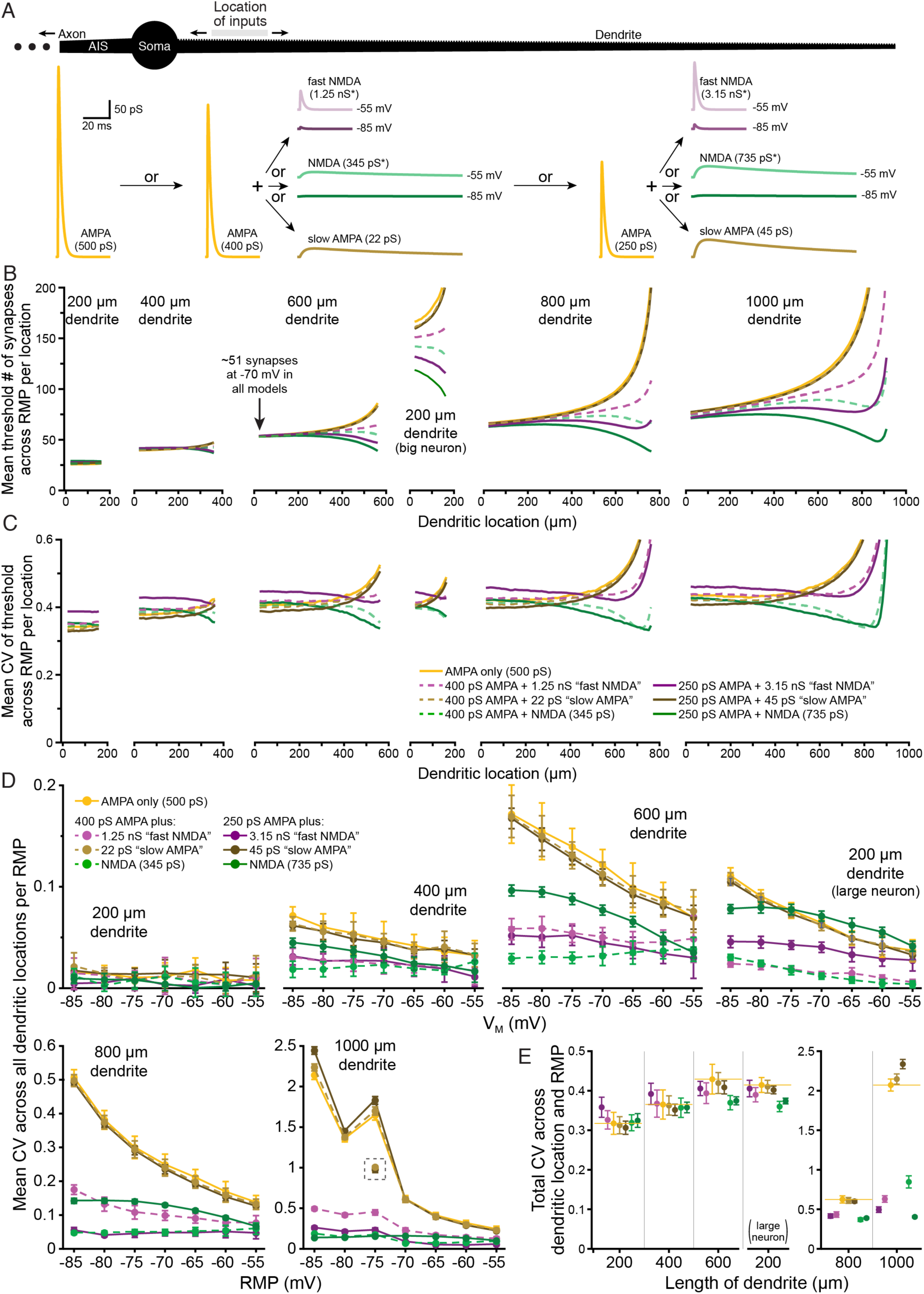
Differential impact of voltage-dependence and kinetics on the fidelity of synaptic integration. ***A***, Diagram of ball-and-stick neuron (top) receiving spatiotemporal patterns of synaptic input to 50-µm spans of dendrite to determine the number of synaptic inputs necessary for action potential generation. Traces below show the synaptic conductances tested: a 500 pS AMPA-only conductance, a 400 pS AMPA conductance “doped” with one of three additional conductances, including a “fast-NMDA” conductance (*nominally 1.25 nS maximal conductance, reaching 4.6 pS at -85 mV, and 49 pS at -55 mV, as shown), a “slow-AMPA” conductance with NMDA kinetics (22 pS), or a nominally 345 pS NMDA conductance (*reaching 1.3 and 13.6 pS at -85 and -55 mV, respectively), or a 250 pS AMPA conductance doped with relatively larger “fast-NMDA”, “slow-AMPA”, and NMDA conductances). All conductances are shown to scale, and were titrated such that each generated a mean threshold of ∼51 synapses in the proximal end of the 600 µm dendrite at -70 mV (arrow in ***B***). ***B***, Plots of mean synaptic thresholds for inputs across all tested RMPs for each 50-µm span of dendrite (incremented at 10-µm intervals) for inputs having the indicated synaptic conductances in dendrites of the indicated lengths (y-axes cut off at 200 synapses; color coding as in panel ***C***). Note that we also tested a 200-µm dendrite attached to a “large neuron” having three additional 600 µm dendrites (middle plot; see also **Figure 3A**). ***C***, Plots of mean CVs for thresholds calculated across seven resting membrane potentials (RMPs; -85 to -55 mV) for each color-coded synaptic conductance at each dendritic location (i.e., measures of voltage-dependent variability of threshold for each given dendritic location) for the different length dendrites (as indicated in ***B***). Y-axes cut off at 0.6. ***D***, Plots of mean CVs (± standard deviations) for synaptic thresholds for each color-coded conductance measured across all dendritic locations within each RMP (i.e., a measure of location-dependent variability in threshold at each RMP). For trials in the 1,000 µm dendrite at -75 mV, the 500 pS AMPA and slow AMPA models hit very high thresholds (>3,000) in their most distal measurable compartment, which generated large CVs at that RMP. CVs calculated for those conductances without the last dendritic location are plotted in the grey dashed box. ***E***, Plots of mean “total CVs” (± standard deviations) calculated across all dendritic locations and all RMPs for neurons with the indicated dendritic lengths and color-coded synaptic conductances.

**Figure 5.**
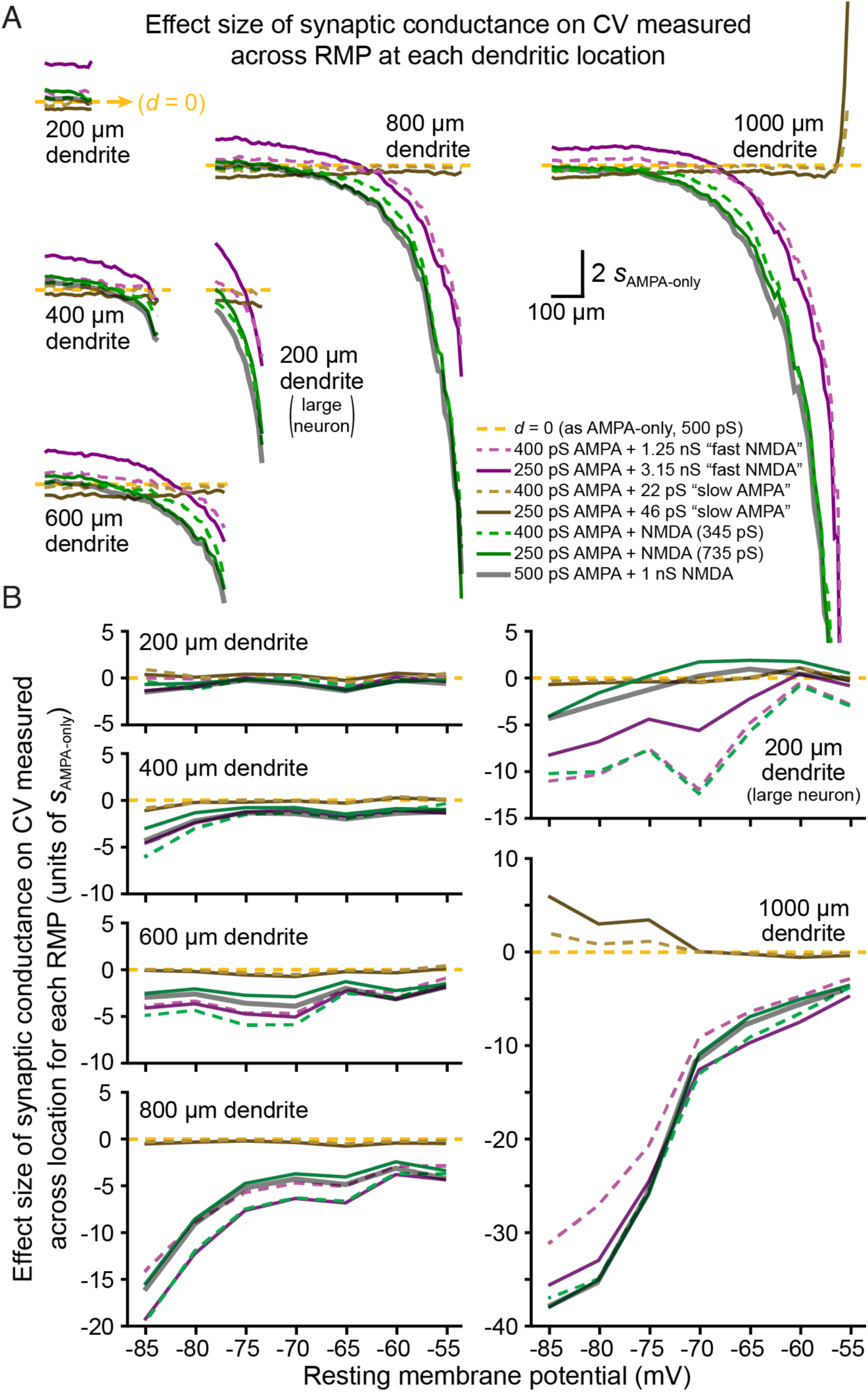
Effect sizes of synaptic conductance on variability of EPSP-spike coupling across RMP and dendritic location. ***A***, Plots of effect sizes of conductance-dependent changes in CV calculated across RMP for each dendritic location (in units of *s*_AMPA-only_, see Methods). Yellow dashed lines indicate *d* = 0 (i.e., identical to 500 pS AMPA-only trials), with effect sizes above that line reflecting larger-than-AMPA-only CVs, while values below the dashed-yellow lines indicate smaller-than-AMPA-only CVs. Most data are from **Figure 4**, and conductance-types are similarly indicated by color. Effect sizes of our normal AMPA (500 pS) conductance combined with 1 nS NMDA (i.e., from **Figure 2C**) are shown as a reference with thick gray semi-transparent lines. Inputs with the “fast-NMDA” conductance tended to generate *more* variability in synaptic thresholds across RMP, especially in short or proximal dendrites, while inputs with slow kinetics exhibited smaller CVs relative to AMPA-only inputs Combining voltage-dependence with slow kinetics (e.g., green and gray “NMDA” traces) reduced variability in synaptic thresholds across RMP at all locations in most dendrites. Plots for 1,000 µm dendrite truncated at -20 *s*_AMPA-only_. ***B***, Plots of effect sizes for changes in CV calculated across all dendritic locations within each RMP. Conductance types are indicated by color, as in ***A***. Voltage-dependent conductances greatly reduced threshold variability across location within a given RMP (an effect magnified at hyperpolarized RMPs), whereas slow kinetics in the absence of voltage-dependence had little effect.

**Figure 6.**
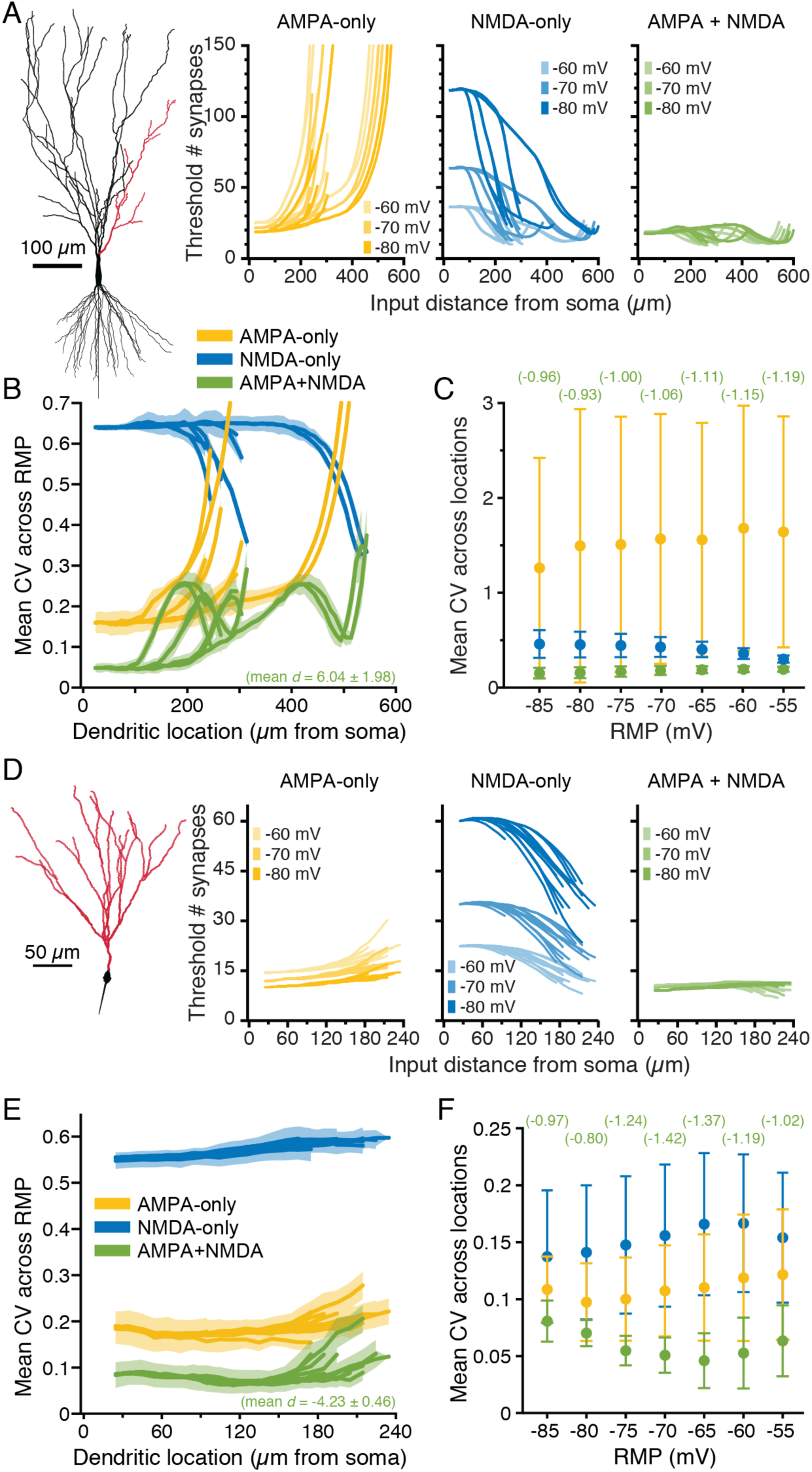
Impact of synaptic conductance on synaptic integration in realistic neuron morphologies. ***A***, Plots of the mean threshold number of synaptic activations necessary to depolarize the soma of a CA3 pyramidal neuron by 5 mV at progressively more distal 50-µm dendritic segments (along red dendrites in diagram at left) experiencing ten static-random spatiotemporal patterns of synaptic input. Colors (yellow, blue, and green) indicate simulations with AMPA, NMDA, and both AMPA and NMDA conductances, respectively, at three different RMPs (−60, -70, and -80 mV). Y-axes are truncated at 200 synapses to see differences in proximal dendrites. ***B***, Plots of mean CVs for synaptic thresholds calculated across seven RMPs (−85 mV to -55 mV) for each dendritic location. Synaptic conductances indicated by color, with shaded regions indicating standard deviations. The mean effect size (*d*, in units of *s*_AMPA-only_) for CVs from AMPA-plus-NMDA simulations (relative to AMPA-only conductance) is shown in green at the bottom of the graph. ***C***, Comparisons of mean (± standard deviation) CVs calculated for synaptic thresholds across dendritic locations within each of seven RMPs. Colors as in ***B***. Effect sizes for AMPA-plus-NMDA trials (vs AMPA-only inputs) are shown in green at top for each RMP (units of *s*_AMPA-only_). ***D***, Plots of the threshold number of synaptic activations necessary to drive a 5 mV somatic depolarization in a dentate granule neuron (red dendrites in diagram to left) at progressively more distal 50-µm segments experiencing ten expanding static-random spatiotemporal patterns of synaptic input. Colors as in ***B. E***, Plots of mean CVs (with standard deviation in shaded regions) of synaptic thresholds across seven RMPs for the different synaptic conductances (indicated by color) for the dentate granule neuron. ***F***, Comparisons of mean CV (± standard deviation) for synaptic thresholds calculated across dendritic location within each of seven RMPs in the granule neuron. Effect sizes of AMPA-plus-NMDA inputs (relative to AMPA-only inputs) are indicated in green at top for each RMP (units of *s*_AMPA-only_).

**Figure 7.**
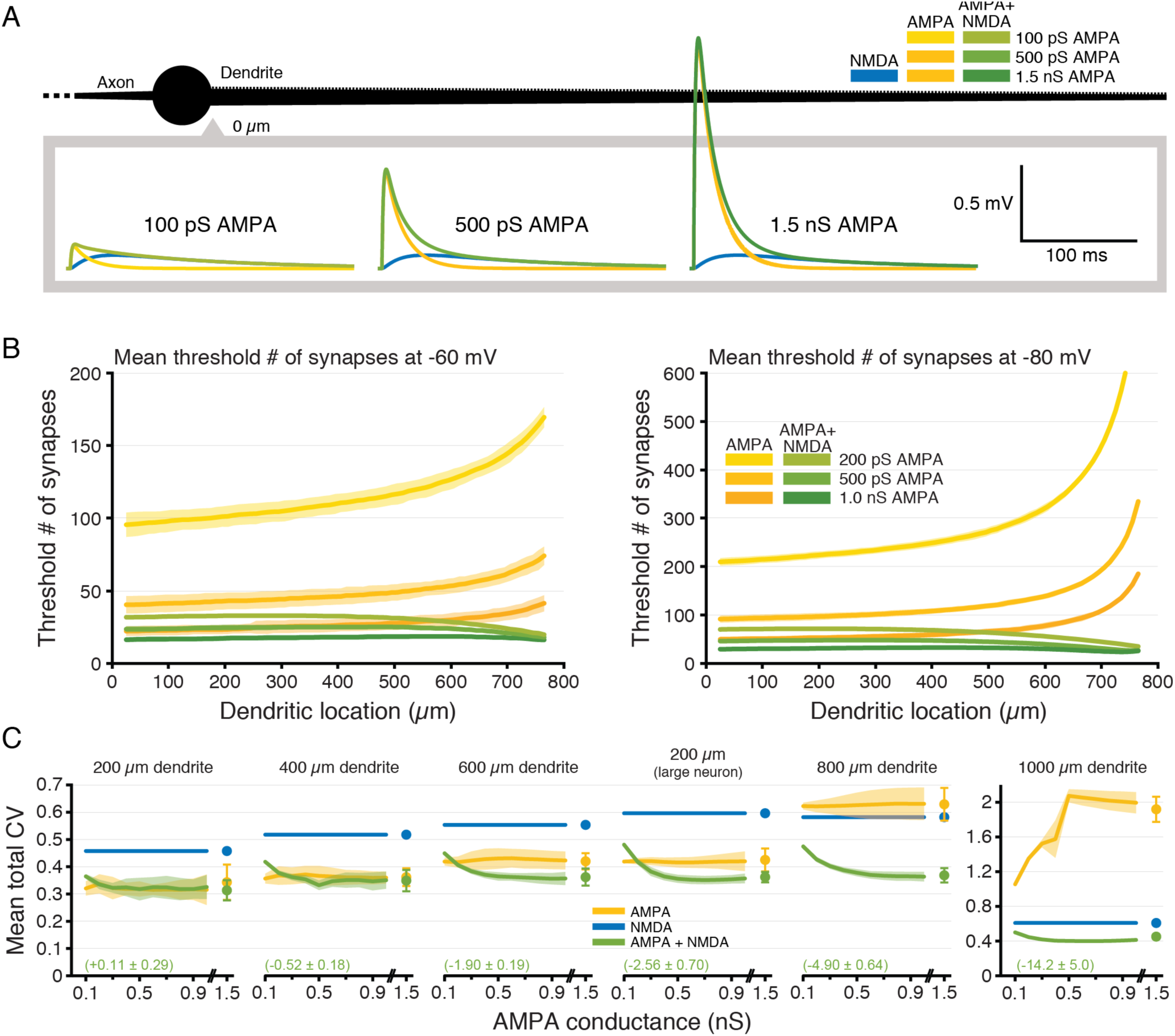
NMDA-dependent stabilization of EPSP-spike coupling occurs over a range of AMPA-to-NMDA conductance ratios. ***A***, Somatic EPSPs generated at -70 mV by proximal inputs having the indicated maximum AMPA conductances (maximal NMDA conductance held steady at 1 nS) in a ball-and-stick neuron (black; not drawn to scale). ***B***, Plots of the mean number of synaptic activations necessary to fire an action potential in an 800-µm-long ball-and-stick neuron resting at -60 mV (left) or -80 mV (right) with peak AMPA conductance set to 0.2, 0.5, or 1 nS, either alone (yellow) or together with a 1 nS NMDA conductance (green). Y-axes for -80 mV data limited to 600 synapses for clarity of differences in proximal locations. ***C***, Plots of mean CVs for thresholds (± standard deviation; shaded regions) for each dendritic length (including the 200 µm dendrite on a “large neuron” with three additional 600-µm dendrites) for inputs having only AMPA (yellow) or AMPA plus NMDA (green) conductances as a function of maximal AMPA conductance (0.1 to 1.5 nS). Mean effect sizes for AMPA+NMDA inputs (relative to AMPA-only inputs, ± standard deviations) calculated for all AMPA conductances ≥ 400 pS are shown in green. As a reference, the CV (± standard deviation) for NMDA-only synapses is shown in blue. Note the Y-axis scale change for the 1,000 µm dendrite.

### Synergistic interaction of voltage-dependence and slow kinetics of the NMDA conductance

The NMDA conductance differs from the AMPA conductance in two key ways: it is voltage-dependent and has slower kinetics. To determine the relative impact of these properties on EPSP-spike coupling, we modified our simulations to measure variability in synaptic thresholds independently across location and RMP (**Figure 4**).

To do this, we started with our AMPA-only model (500 pS) and then replaced a portion of that conductance (100 or 250 pS) with either a) the normal NMDA conductance, b) a conductance having NMDA-like voltage-dependence with AMPA-like kinetics (“fast-NMDA”), or c) a conductance with NMDA-like kinetics but lacking voltage-dependence (“slow-AMPA”). These added conductances were titrated so that each model generated a similar synaptic threshold for inputs to the proximal end of the 600 µm dendrite at -70 mV (∼ 51 synapses). The resulting synaptic conductances are shown to scale in **Figure 4A** following activation of individual synapses under voltage-clamp (voltage-dependent conductances shown for activations at -85 and -55 mV, as indicated). As in **Figure 2**, for each dendritic location (50-µm spans, incremented at 10-µm intervals), and RMP (from -85 to -55 mV, in 5-mV increments), we measured synaptic thresholds for action potential generation in response to each of ten randomized spatiotemporal patterns of synaptic input (**Figure 4B**). We then calculated the CV of synaptic thresholds across RMPs for each dendritic location (**Figure 4C**), across all dendritic locations at a given RMP (**Figure 4D**), and across all dendritic locations and RMPs (“total CV”; **Figure 4E**). This allowed us to dissociate the relative impacts of voltage-dependance and conductance kinetics on the fidelity of EPSP-spike coupling. Adding back an AMPA conductance with slower kinetics (the brown plots from simulations including the “slow-AMPA” conductance in **Figure 4C**) modestly reduced the CV of synaptic thresholds measured across RMP at a given dendritic location, especially in shorter dendrites or in proximal locations in longer dendrites. On the other hand, adding NMDA-like voltage-dependence with fast, AMPA-like, kinetics (“fast-NMDA”; purple plots in **Figure 4C**) increased the CV of synaptic thresholds measured across RMP at individual dendritic locations in small and/or proximal dendrites. Both of these effects were “dose-dependent,” in that they were larger when they contributed proportionally more to the total synaptic conductance (compare dashed vs solid lines in **Figure 4C**).

When our normal NMDA conductance (titrated, as above) was added to these smaller AMPA conductances (green traces in **Figure 4C**), CVs for synaptic thresholds measured across RMP were intermediate to those generated by fast-NMDA and slow-AMPA conductances, being only marginally larger than for AMPA-only inputs in short dendrites (or at proximal dendritic locations in longer dendrites) but becoming progressively lower at more distal dendritic locations. Overall, mean CVs of synaptic thresholds measured across RMP at all dendritic locations with titrated N M D A conductances were lower than for AMPA-only inputs in all dendrites longer than 200 um, and in the the 200-µm dendrite attached to a larger neuron with three additional 600-µm dendrites (effect sizes ranging from -0.26 *s*_AMPA-only_ in the 400-µm dendrite to -1.91 *s*_AMPA-only_ in the 200-µm dendrite attached to a larger neuron). Effect sizes of conductance-dependent changes in synaptic threshold variability across RMP are plotted for each dendritic location in **Figure 5A** (below).

The impact of synaptic conductance on synaptic threshold variability was quite different when measured across all dendritic locations within a given RMP (**Figure 4D**). For all dendrites greater than 200 µm (including in the 200-µm dendrite attached to a larger neuron), the presence of the “fast-NMDA” conductance greatly lowered the CV of synaptic thresholds within any given RMP, while adding the “slow-AMPA” conductance had little impact on synaptic threshold variability across dendritic locations. The combination of voltage-dependence and slow kinetics (i.e., our titrated standard NMDA conductance; green traces) generally lowed CV for synaptic thresholds measured across dendritic locations for a given RMP. For both fast-NMDA and regular NMDA conductances, the dose-dependency was non-linear, as the smaller dose typically generated the largest reductions in CV (relative to AMPA-only conductance; **Figure 4D**). The effect sizes of these synaptic conductances on threshold variability across location within a given RMP (relative to 500 pS AMPA alone) are shown in **Figure 5B**.

Finally, we measured “total CV” of synaptic thresholds across all dendritic locations and all RMPs for ten patterns of synaptic input (**Figure 4E**). In smaller neurons (those with 200-µm and 400-µm dendrites), adding the “fast-NMDA” conductance (purple symbols) slightly increased CV relative to AMPA conductance alone. On the other hand, adding “slow-AMPA” kinetics (brown symbols) modestly reduced CV relative to AMPA-only inputs in most models. These effects were reversed in larger neurons such that the “slow-AMPA” conductance increased mean total CV in the 1,000-µm dendrite, while the voltage-dependent “fast-NMDA” conductance reduced total CV for all dendrites longer than 400 µm, and in the 200-µm dendrite on a larger neuron. This likely reflects the ability of the “fast-NMDA” conductance to effectively lower CV across distance in larger neurons by avoiding or delaying the exponential rise in synaptic thresholds observed for AMPA-only inputs (**Figure 4C**), and that slower kinetics (i.e., “slow-AMPA” models) have reduced efficacy in lowering CV across RMPs at distal dendritic locations (**Figure 4C**).

Combining voltage-dependence with slow kinetics (i.e., the titrated normal NMDA conductance; green symbols in **Figure 4E**) led to overall lower total CVs for synaptic thresholds in all dendrites longer than 200 µm, including in the 200-µm dendrite attached to a larger neuron. Mean total CVs (across 10 synaptic input patterns) and effect sizes for conductance-dependent changes in total CV (relative to the 500 pS AMPA-only conductance) are shown in **Table 2**. In most dendrites, combining voltage-dependence with slower kinetics had a synergistic effect in lowering total CV of synaptic thresholds to a greater extent than either manipulation on its own, often with “large” effect sizes (i.e. > 1 *s*_AMPA-only_).

**Table 2:** Effect sizes for Figure 4E

| Dendritic length (μm) | Conductance | gAMPA | Mean "total" CV | SD | Effect size (d) | gAMPA | Mean "total" CV | SD | Effect size (d) |
| --- | --- | --- | --- | --- | --- | --- | --- | --- | --- |
| 200 | AMPA-only | 500 | 0.317 | 0.048 |  |  |  |  |  |
|  | AMPA + fast NMDA | 400 | 0.326 | 0.023 | 0.325 | 250 | 0.358 | 0.025 | 1.469 |
|  | AMPA + slow AMPA | 400 | 0.313 | 0.022 | -0.169 | 250 | 0.306 | 0.017 | -0.394 |
|  | AMPA + NMDA | 400 | 0.319 | 0.020 | 0.057 | 250 | 0.325 | 0.017 | 0.263 |
| 400 | AMPA-only | 500 | 0.365 | 0.280 |  |  |  |  |  |
|  | AMPA + fast NMDA | 400 | 0.367 | 0.034 | 0.060 | 250 | 0.392 | 0.027 | 0.721 |
|  | AMPA + slow AMPA | 400 | 0.363 | 0.024 | -0.071 | 250 | 0.352 | 0.014 | -0.371 |
|  | AMPA + NMDA | 400 | 0.357 | 0.024 | -0.223 | 250 | 0.358 | 0.013 | -0.205 |
| 600 | AMPA-only | 500 | 0.429 | 0.038 |  |  |  |  |  |
|  | AMPA + fast NMDA | 400 | 0.394 | 0.026 | -0.940 | 250 | 0.406 | 0.018 | -0.623 |
|  | AMPA + slow AMPA | 400 | 0.420 | 0.027 | -0.253 | 250 | 0.409 | 0.016 | -0.546 |
|  | AMPA + NMDA | 400 | 0.370 | 0.018 | -1.578 | 250 | 0.375 | 0.010 | -1.450 |
| 200 (on large neuron) | AMPA-only | 500 | 0.414 | 0.019 |  |  |  |  |  |
|  | AMPA + fast NMDA | 400 | 0.390 | 0.018 | -1.308 | 250 | 0.405 | 0.021 | -0.476 |
|  | AMPA + slow AMPA | 400 | 0.410 | 0.016 | -0.222 | 250 | 0.402 | 0.010 | -0.666 |
|  | AMPA + NMDA | 400 | 0.360 | 0.013 | -2.886 | 250 | 0.374 | 0.007 | -2.140 |
| 800 | AMPA-only | 500 | 0.625 | 0.046 |  |  |  |  |  |
|  | AMPA + fast NMDA | 400 | 0.434 | 0.035 | -4.131 | 250 | 0.415 | 0.027 | -4.535 |
|  | AMPA + slow AMPA | 400 | 0.612 | 0.037 | -0.283 | 250 | 0.601 | 0.020 | -0.520 |
|  | AMPA + NMDA | 400 | 0.369 | 0.023 | -5.537 | 250 | 0.389 | 0.010 | -5.104 |
| 1000 | AMPA-only | 500 | 2.071 | 0.076 |  |  |  |  |  |
|  | AMPA + fast NMDA | 400 | 0.630 | 0.041 | -18.964 | 250 | 0.495 | 0.036 | -20.737 |
|  | AMPA + slow AMPA | 400 | 2.147 | 0.065 | 1.002 | 250 | 2.337 | 0.058 | 3.493 |
|  | AMPA + NMDA | 400 | 0.847 | 0.077 | -16.111 | 250 | 0.405 | 0.010 | -21.928 |

To further compare the impact of synaptic increase in the CVs for synaptic thresholds across RMPs in shorter dendrites and in the proximal two-thirds of longer dendrites (those ≥ 600 µm), with effect sizes for the larger dose of “fast-NMDA” (3.15 nS, paired with 250 pS AMPA) averaging +1.08 ± 0.26 *s*_AMPA-only_ (i.e., more variable thresholds across RMP relative to AMPA-only inputs) in shorter (200 µm and 400 µm) dendrite models and in the proximal two-thirds of longer dendrites. At distal locations in longer dendrites, effect sizes for NMDA-containing models became negative (i.e., reduced variability across RMP) and very large relative to AMPA-only inputs (**Figure 5**). This occurred because adding a voltage-dependent component (regardless of kinetics) avoided, or at least delayed, the distance-dependent exponential climb in synaptic thresholds exhibited at distal dendritic locations (see **Figure 4B**) and the corresponding increases in synaptic threshold variability (see **Figure 4C**).

Adding the larger dose of “slow-AMPA” conductance moderately lowered variability of synaptic thresholds across RMPs for most proximal dendritic locations, with effect sizes averaging -0.46 ± 0.13 *s*_AMPA-only_ in all models > 200 µm, including in the 200 µm dendrite conductance on the variability of EPSP-spike coupling, as measured across all RMPs (for a given dendritic location) or across all dendritic locations (for a given RMP), we plotted the effect sizes for our various synaptic conductances (relative to threshold variability observed for AMPA-only inputs) in **Figure 5**. This essentially transforms data in **Figures 4C** and **4D** into measures of effect size (in units of *s*_AMPA-only_) to allow comparison of the differential impact of voltage-dependence and slow kinetics on synaptic threshold variability across RMPs (**Figure 5A**) or across all dendritic locations (**Figure 5B**). Included in **Figure 5** are results f rom our normal AMPA + NMDA conductance (500 pS AMPA + 1 nS NMDA; thick semi-transparent gray lines), which tended to track closely to the titrated NMDA conductances (green lines). As shown in **Figure 5A**, adding the “fast-NMDA” conductance induced a dose-dependent attached to a larger neuron (**Figure 5A**).

Combining voltage-dependence with slow kinetics (e.g., green “NMDA” traces in **Figures 4C** and **5A**) revealed a location-dependent reduction in variability of synaptic thresholds measured across RMPs that was similar to that observed when adding the “fast NMDA” conductance (compare green and purple traces in **Figure 5A**). However, relative to fast NMDA trials, the slower kinetics of the normal NMDA conductance led consistently to lower CVs for synaptic thresholds at all dendritic locations in all model neurons. Mean effect sizes for all dendritic locations ranged from negligible (+0.06 *s*_AMPA-only_ in the 200-µm dendrite) to very large (−5.2 in the 1,000-µm dendrite), and always became larger with increasing distance from the soma. As with the “fast-NMDA” conductance, the enormous effect sizes in distal dendrites reflects primarily the earlier (i.e., at more proximal dendritic locations) exponential increases in synaptic thresholds in AMPA-only models, rather than the more modest distance-dependent reduction in CV occurring in the presence of the NMDA conductance (**Figure 4C**).

The impact of synaptic conductance on synaptic threshold variability measured across location within a given RMP is shown in **Figure 5B**. Regardless of kinetics, the addition of voltage-dependence greatly reduced the CV of synaptic thresholds across location (**Figure 5B**), and this effect was magnified in larger dendrites and at more hyperpolarized RMPs. The presence of slow kinetics in the absence of voltage-dependence had little impact on synaptic threshold variability across dendritic location, except in the longest dendrite (1,000 µm) at hyperpolarized RMPs, where variability increased relative to AMPA-only inputs.

Together, these results indicate that the impact of the NMDA conductance on the variability of synaptic thresholds across dendritic location and RMP reflects a synergistic interaction of multiple properties of the NMDA conductance, with its voltage-dependence primarily enhancing fidelity across dendritic location within any given RMP, and its slower kinetics tending to enhance fidelity across RMP at a given dendritic location. These properties act together to greatly reduce the variability of EPSP-spike coupling across both RMP and dendritic location, relative to AMPA-only synaptic inputs.

### Synaptic integration in morphologically realistic neurons

The simulations above reveal that, in simplified neuron morphologies, the presence of both AMPA and NMDA conductances at synapses reduces location- and RMP-dependent variability of somatic responses to spatially restricted barrages of synaptic input. To determine the impact of synaptic conductance on synaptic integration in realistic neuronal morphologies, we placed spinous synapses at 1-µm intervals across the dendritic trees of two reconstructed neurons: a relatively large CA3 pyramidal neuron (**Figure 6A**), and a smaller dentate granule neuron (**Figure 6D**). Because of their large dendritic arbors, it was not possible to drive action potential generation in these neurons with spatially restricted (50 µm) synaptic barrages, as even extreme depolarization of distal dendrites (e.g., clamping 50-µm spans to 0 mV) failed to depolarize the axon initial segment to action potential threshold. Instead, we set arbitrary thresholds of somatic depolarization (3 mV above RMP for the CA3 neuron, and 5 mV above RMP for the smaller dentate granule neuron) and iteratively activated expanding stochastic patterns of synaptic input at different dendritic locations until somatic voltage thresholds were realized. RMPs were nominally set between -85 and -55 mV, in 5 mV increments, and synaptic barrages delivered to 50-µm spans along the dendritic branches indicated in red in **Figures 6A** and **D** (at 10-µm increments). In the CA3 neuron, the threshold number of AMPA-only inputs necessary to generate a 3 mV somatic depolarization increased with distance from the soma, or with depolarization of the RMP, as voltage attenuation and reduced driving force impaired synaptic depolarization of the soma (yellow plots in **Figure 6A**). In contrast, when synapses contained only the NMDA conductance, the threshold number of synapses to depolarize the soma by 3 mV decreased with distance from the soma and/or with depolarization of the RMP, as distance-dependent increases in local input impedance, and/or depolarization of the RMP, enhanced the voltage-dependent NMDA conductance (blue plots in **Figure 6A**). Combining both AMPA and NMDA conductances generated threshold numbers of synapses that were generally lower and more uniform across dendritic location and RMP (**Figure 6B**, green plots). Whether measured across RMP (for a given dendritic location; **Figure 6B**) or across dendritic location (within a given RMP; **Figure 6C**), CVs were always less when AMPA and NMDA conductances were combined. Across RMPs, CVs of AMPA+NMDA inputs were reduced, on average, by 6.04 ± 1.98 *s*_AMPA-only_ (**Figure 6B**), while effect sizes for reductions in CV across dendritic location (within a given RMP) ranged from -0.93 to -1.19, primarily because of large variability in the CVs of AMPA-only trials (compare the standard deviations of AMPA-only and AMPA+NMDA trials in **Figure 6B**).

Across all dendritic locations and RMPs, total CVs in AMPA-only trials averaged 1.83 ± 0.10 across ten patterns of synaptic input, but were only 0.19 ± 0.01) in AMPA+NMDA trials, with an effect size of -16.6 *s*_AMPA-only_ for AMPA+NMDA inputs relative to AMPA-only trials. These results demonstrate that the combined presence of synaptic AMPA and NMDA conductances greatly reduces distance- and RMP-dependent variability of synaptic integration (relative to AMPA-only inputs) in a realistic neuron morphology.

Similar results were observed in the smaller dentate granule neuron (**Figure 6D**), where the threshold number of AMPA-mediated synaptic inputs increased with distance from the soma, or with depolarization of the RMP (**Figure 6D**, yellow plots), due to distance-dependent voltage attenuation and reduced synaptic driving force, respectively. Conversely, for NMDA-mediated inputs, synaptic thresholds decreased with distance from the soma, or with depolarization of the RMP (**Figure 6D**, blue plots). As was true in the CA3 neuron, the combined presence of synaptic AMPA and NMDA conductances in the dentate granule neuron minimized the impact of dendritic location and/or RMP on synaptic thresholds (**Figure 6D**, green plots) and greatly reduced the CVs of thresholds calculated across RMPs for increasingly distant dendritic locations (**Figure 6E**) or across dendritic location within a given RMP (**Figure 6F**). Mean total CVs for the ten spatiotemporal input patterns, calculated across all dendritic locations and RMPs, were 0.19 ± 0.02, 0.52 ± 0.01, and 0.08 ± 0.01, for AMPA-only, NMDA-only, and AMPA-and-NMDA conductances, respectively, with an effect size of -5.18 *s*_AMPA-only_ for AMPA+NMDA inputs relative to AMPA-only inputs.

### Impact of AMPA-to-NMDA conductance ratios on synaptic integration

The simulations described above reveal that combining AMPA and NMDA conductances can increase the fidelity of synaptic integration, as measured at the soma and axon, across dendritic locations and/or RMPs. However, while the number of synaptic NMDA receptors is fairly uniform across most synapses (on the order of 10 receptors per synapse; Racca et al., 2000; Nimchinsky et al., 2004; Noguchi et al., 2005), synaptic AMPA conductances can range from zero (i.e., in “silent” synapses) to >1.5 nS (Beaulieu-Laroche and Harnett, 2018), and are dynamic in response to synaptic plasticity (for review, see Herring and Nicoll, 2016). Thus, the ratio of AMPA-to-NMDA conductance at synapses is variable. To determine the impact of AMPA-to-NMDA ratio on the fidelity of synaptic integration, we varied the maximal AMPA conductance from 0.1 to 1.5 nS while keeping the maximal NMDA conductance steady at 1 nS, and measured the minimum number of synaptic activations necessary to trigger an action potential in ball-and-stick neurons of various dendritic lengths (**Figure 7**). Regardless of AMPA conductance magnitude, the threshold number of synaptic inputs for action potential initiation increased sharply with distance from the soma for AMPA-only inputs, while thresholds for inputs containing both AMPA and NMDA conductances were relatively independent of distance (**Figure 7B**). In long dendrites (> 600 µm) the mean CV for the threshold number of inputs was lowest when both conductances were present, regardless of AMPA-to-NMDA ratio (**Figure 7C**). In ball-and-stick model neurons with short dendrites, when AMPA conductances were small, adding an NMDA conductance did not significantly reduce threshold CVs, as the NMDA conductance became the dominant driver of somatic depolarization, and CVs converged toward those of NMDA-only inputs. This reflects the tight electrotonic structure of small model neurons, as adding NMDA conductance reduced the CV of EPSP-spike coupling across most AMPA-NMDA ratios in the 200-µm dendrite when it was attached to a larger neuron with three additional 600-µm dendrites (“large neuron” in **Figure 7C**; compare also results from the short-but-branched dendritic arbor of the dentate granule neuron in **Figure 6D-F**). For all AMPA conductances ≥ 400 pS, the mean effect size of adding NMDA conductance (relative to AMPA-only inputs) varied between negligible (*d* = +0.11 ± 0.29 *s*_AMPAonly_ in the 200 µm dendrite) to enormous (*d* = -15.2 ± 5.0 *s*_AMPAonly_ in the 1,000 µm dendrite), with mean effect sizes always being greater than 1.0 for dendrites ≥ 600 µm (including in the 200-µm dendrite attached to a larger neuron; **Figure 7C**). Thus, the voltage-dependence of the NMDA conductance enhances the fidelity of synaptic integration across a wide range of physiologically relevant AMPA-to-NMDA ratios.

## Discussion

Synaptic integration, the process by which patterns of synaptic input are transduced into action potential initiation (also known as “EPSP-spike coupling”), is the core of neural computation. We tested the impact of two prominent and often co-expressed glutamate receptor subtypes, AMPA and NMDA receptors, on the fidelity of synaptic integration across dendritic location and neuronal state (i.e., initial membrane potential). Our results demonstrate that the kinetics and voltage-dependence of the NMDA conductance act synergistically to counterbalance the impact of distance-dependent signal attenuation and the reduced synaptic driving forces occurring at depolarized membrane potentials, effectively increasing the fidelity of EPSP-spike coupling across both spatial and voltage domains (as summarized in **Figure 8**). This intrinsic consequence of combining synaptic AMPA and NMDA receptors occurs over a broad range of neuron morphologies and AMPA-to-NMDA conductance ratios, and is independent of, but fully compatible with, its well characterized role in gating synaptic plasticity.

**Figure 8.**
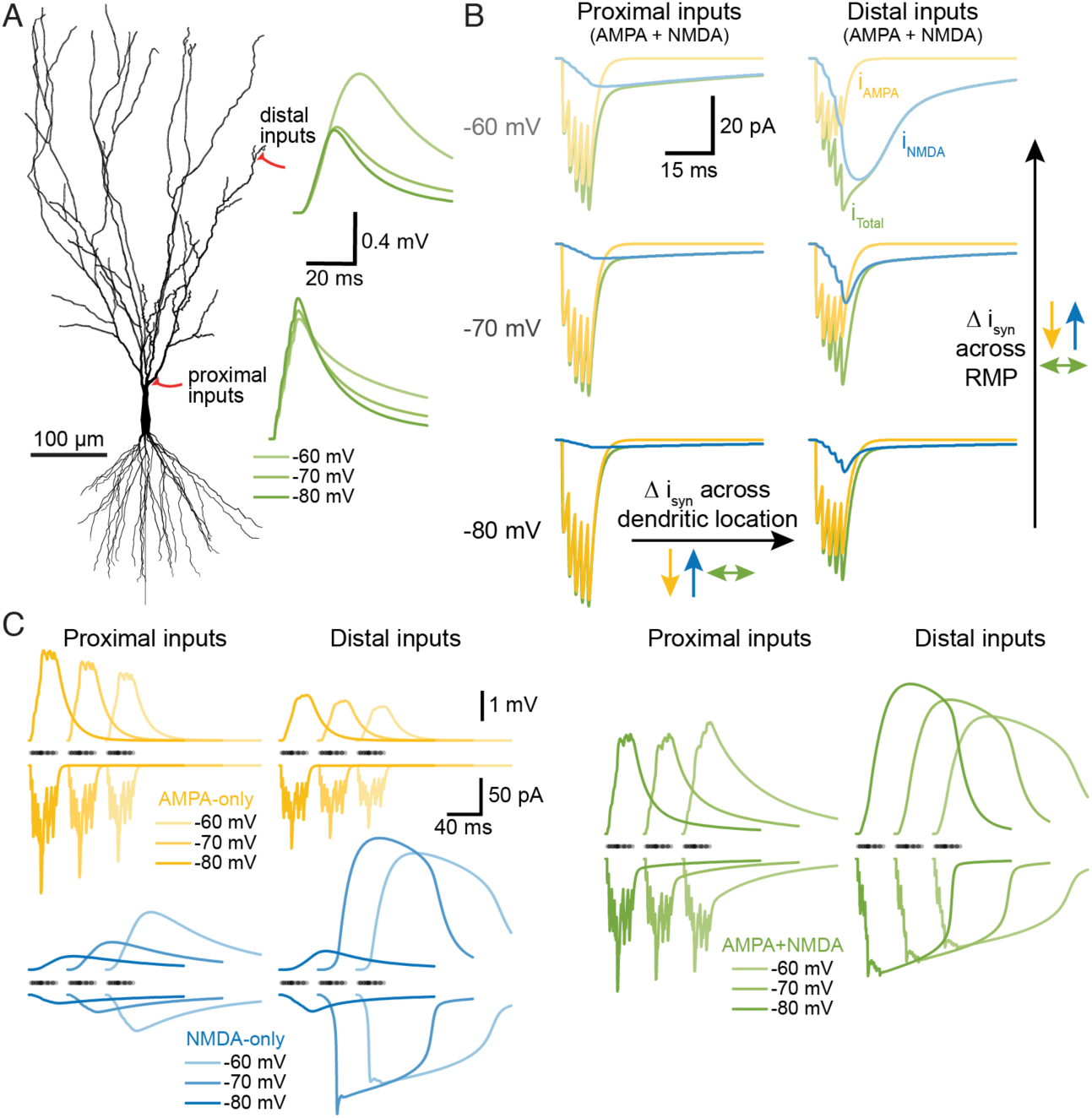
Summary of NMDA-dependent stabilization of synaptic integration across dendritic location and RMP. ***A***, Morphology of the CA3 neuron indicating the locations of proximal and distal synaptic inputs to the apical dendrite. Traces at right show the somatic responses to sequential activation of five AMPA+NMDA synapses at neighboring spines (at 2 ms intervals, distal to proximal) at three different RMPs (as indicated by color-depth) for each input location (distal vs proximal) in a fully passive model. ***B***, Synaptic currents (i_syn_) generated by activation of the five proximal (left) or five distal (right) inputs at three RMPs. Total synaptic currents (i_Total_) are shown in green, with AMPA (i_AMPA_; yellow) and NMDA (i_NMDA_; blue) components superimposed. Notice that AMPA currents are reduced (yellow arrows), while NMDA currents are enhanced (blue arrows), as inputs are moved from proximal to distal locations, or as RMP is depolarized from -80 mV to -60 mV. These opposing effects of location and RMP limit the variability of the total synaptic current across location and RMP (green arrows). ***C***, Similar to ***B***, but showing somatic voltage responses (upper traces) and summed synaptic currents (lower traces) for a barrage of 25 synaptic inputs with stochastic timings and spine locations for AMPA-only (yellow), NMDA-only (blue) or AMPA+NMDA (green) inputs. Timings of synapses indicated with semi-translucent dots above current traces. Note how AMPA-only currents and somatic EPSPs (yellow) get smaller with depolarization or distance from the soma, while NMDA-only currents and EPSPs (blue) get larger with distance and depolarization. Combining AMPA and NMDA conductance (green) leads to less variation in the amplitudes of both synaptic current and somatic EPSPs across RMPs and/or dendritic locations.

### Interaction of electrotonic structure and synaptic conductance

EPSPs are shaped by the electrotonic structure of the dendritic tree; in narrow distal dendrites, small surface areas limit local membrane conductance and capacitance, yielding greater input impedance and larger and faster local EPSPs in response to synaptic currents (Gulledge et al., 2005). This can reduce synaptic currents in distal dendrites, as greater local depolarization during EPSPs limits synaptic driving forces at the peak of the EPSP. At the same time, dendrites act as low-pass filters for EPSPs spreading from the site of synaptic input toward the soma. Thus, in the absence of voltage-dependent conductances, the somatic efficacy of synaptic input diminishes with distance from the soma, thereby requiring a greater number of synaptic inputs to reach action potential threshold. Our results with AMPA-only synaptic inputs are fully consistent with these well-described effects of dendritic cable properties (e.g., **Figures 1** and **2**). NMDA conductances, on the other hand, generate larger synaptic currents in distal, high impedance dendritic compartments due to their enhanced conductance at depolarized membrane potentials (e.g., Gulledge et al., 2012; Lajeunesse et al., 2013). Our main result is that the combined presence of AMPA and NMDA receptors intrinsically lowers location- and RMP-dependent variability in synaptic efficacy by boosting synaptic conductance preferentially in high-impedance dendritic compartments (Branco and Hausser, 2011; Gulledge et al., 2012) and/or when neurons are in depolarized states (Branco et al., 2010), conditions in which synaptic driving forces are diminished (see **Figures 2** and **8**). NMDA-dependent stabilization of synaptic integration was robust across neuron morphologies (**Figures 2, 3**, and **6**) and over a wide range of AMPA-to-NMDA conductance ratios (**Figure 7**), and persisted in the presence of active properties in dendrites (**Figure 3C**) that can contribute to voltage-dependent amplification of inward current following synaptic activation (Magee and Johnston, 1995; Lipowsky et al., 1996; Branco et al., 2010).

By doping the AMPA conductance with voltage-dependence (“fast-NMDA”) or slower, non-voltage-dependent kinetics (“slow-AMPA”; see **Figures 4** and **5**), we were able to measure the relative impact of these two distinct properties of the NMDA conductance. Adding voltage-dependence to the AMPA conductance reduced variability of synaptic thresholds for action potential generation when measured across dendritic location within a given RMP, especially in long dendrites, but generally increased variability of synaptic thresholds when measured across RMP at a given dendritic location. This reflects the gradual recruitment of the NMDA conductance that occurs as synaptic barrages occur progressively in ever-more-distal higher-impedance locations that leads to larger distal synaptic currents (see **Figures 2B** and **8**). On the other hand, the increased variability of EPSP-spike coupling when barrages were initiated at different RMPs in proximal dendrites reflects the non-linear amplification of synaptic responses with increased depolarization, leading to a greater range of synaptic currents across voltage at any given location.

The slower kinetics of NMDA receptors, on their own, reduced variability of synaptic thresholds selectively across RMP at proximal dendritic locations by increasing the opportunity for temporal summation during barrages. At any point within the stochastic barrage, the next iterative input is more likely to summate to a suprathreshold response if EPSP decay is slow. Since the decay of EPSPs depends on the local time constant, which itself is location dependent (i.e., larger near the soma; compare, for instance, the decay rates of proximal vs distal EPSPs having identical kinetics in **Figure 1A**), the impact of slower kinetics will be largest in proximal dendritic locations (see **Figures 4** and **5**). The overall impact of adding voltage-dependent and slower NMDA conductances to AMPA synapse is to greatly reduce the variability of somatic responses and EPSP-spike coupling across both dendritic location and membrane potential state (e.g., **Figures 2** and **6**).

### Impact of NMDA conductance on synaptic integration and network performance

Neurons employ a variety of mechanisms to combat location-dependent variability in synaptic efficacy. In the apical dendrites of CA1 pyramidal neurons, synaptic conductance is scaled in proportion to distance from the soma to minimize location-dependent variability of axonal drive (Magee and Cook, 2000; Nicholson et al., 2006; Shipman et al., 2013). Alternatively, dendrites may express dendritic voltage-gated sodium and calcium conductances that, with sufficient local depolarization, can generate dendritic action potentials that amplify somatic EPSPs (e.g., Magee et al., 1995; Magee and Johnston, 1995; Schiller et al., 1997; Golding and Spruston, 1998; Golding et al., 1999; Milojkovic et al., 2005). Similarly, synaptic NMDA receptors, when activated in sufficient numbers, generate self-sustaining plateau potentials (“NMDA spikes”) that lead to supralinear synaptic summation and larger somatic EPSPs (Schiller et al., 2000; Wei et al., 2001; Polsky et al., 2004; Nevian et al., 2007; Major et al., 2008; Larkum et al., 2009; Branco et al., 2010; Branco and Hausser, 2011; Harnett et al., 2012). These voltage-dependent synaptic and dendritic mechanisms act preferentially in distal dendritic compartments, where input impedance is highest, and facilitate synaptic depolarization of the soma and axon in spite of distance-dependent voltage-attenuation within the dendritic arbor. However, unlike action potentials, NMDA spikes are not all-or-none. Instead, they vary in amplitude and duration in proportion to the number of activated synapses, the local input impedance at the synapse, and the initial membrane potential (Major et al., 2008; Branco et al., 2010; Branco and Hausser, 2011; Gulledge et al., 2012; Farinella et al., 2014). Thus, although highly nonlinear across voltage, NMDA spikes are effectively “graded” in amplitude and duration across dendritic locations (see **Figure 2B**), allowing the NMDA conductance to enhance synaptic current in proportion to electrotonic distance from the soma and axon. Similarly, by opposing the impact of reduced synaptic driving force on AMPA-mediated currents, voltage-dependent NMDA conductances can reduce the variability of total synaptic current during temporal integration of clustered excitatory inputs (Connelly et al., 2016; see also **Figure 8C**).

Over the past several decades, there has been growing appreciation that, in addition to gating many forms of synaptic plasticity, NMDA receptors play an integral role in normal synaptic integration (for reviews, see Hausser and Mel, 2003; Antic et al., 2010; Major et al., 2013). In the distal apical tufts of layer 5 neurons, where high input impedance favors amplification of local EPSPs, NMDA activation boosts synaptic currents such that they can more reliably trigger calcium spikes at an electrically excitable zone at the base of the tuft (Larkum et al., 2009). Indeed, simulations by Larkum et al. (2009) found the threshold number of synapses necessary for initiating an apical trunk calcium spike to be relatively consistent across tuft locations when NMDA conductances were present, but that thresholds for AMPA-only inputs grew exponentially, and to non-physiological synaptic densities, with distance from the initiation zone in the apical dendrite. Thus, the impact of NMDA receptors on the fidelity of EPSP-spike coupling may not be limited to action potentials initiated at the axon, but likely applies more generally for spike initiation at any specialized trigger zone, so long as it is electrotonically downstream (e.g., in a larger compartment) from the site of synaptic input.

### Conclusions

Transduction of synaptic events into patterns of action potential output is the most fundamental neuronal task. It is a process influenced by the strength and kinetics of individual synaptic conductances, the spatiotemporal relationships among them, and the electrotonic properties of the neuron. Nonlinear amplification of synaptic input via NMDA spikes is proposed to increase the “computational power” of neurons (e.g., Wei et al., 2001; Poirazi et al., 2003; Branco and Hausser, 2011). Our results demonstrate that NMDA receptors, via their intrinsic kinetics and voltage-dependence, also provide the advantage of “computational stability” by enhancing the fidelity of EPSP-spike coupling across dendritic domains and membrane potential states. While speculative, it is interesting to consider whether this consequence of NMDA receptor expression may have provided advantages to primitive nervous systems (e.g., in cnidaria; Anctil, 2009; Pierobon, 2012) independent of their role in associative plasticity, which may account for their ancient evolutionary origin in the common ancestor of all metazoans (Ramos-Vicente et al., 2018).

## Author Contributions

A.T.G. conceived the project and wrote the paper. C.L. and A.T.G designed and performed experimental work, analyzed data, and reviewed and edited the paper.

## Acknowledgements

The authors thank Arielle Baker, Nik Dembrow, and Greg Stuart for comments and suggestions on a draft of this manuscript, and Saiko Ikeda for assistance with data analyses.

